# Exploiting deep transfer learning for the prediction of functional noncoding variants using genomic sequence

**DOI:** 10.1101/2022.03.19.484983

**Authors:** Li Chen, Ye Wang

## Abstract

**Motivation:** Though genome-wide association studies have identified tens of thousands of variants associated with complex traits and most of them fall within the noncoding regions, they may not the causal ones. The development of high-throughput functional assays leads to the discovery of experimental validated noncoding functional variants. However, these validated variants are rare due to technical difficulty and financial cost. The small sample size of validated variants makes it less reliable to develop a supervised machine learning model for achieving a whole genome-wide prediction of noncoding causal variants.

**Results:** We will exploit a deep transfer learning model, which is based on convolutional neural network, to improve the prediction for functional noncoding variants. To address the challenge of small sample size, the transfer learning model leverages both large-scale generic functional noncoding variants to improve the learning of low-level features and context-specific functional noncoding variants to learn high-level features toward the context-specific prediction task. By evaluating the deep transfer learning model on three MPRA datasets and 16 GWAS datasets, we demonstrate that the proposed model outperforms deep learning models without pretraining or retraining. In addition, the deep transfer learning model outperforms 18 existing computational methods in both MPRA and GWAS datasets.

**Availability:** https://github.com/lichen-lab/TLVar

**Supplementary Information:** Supplementary data are available at Bioinformatics online.

## 1 INTRODUCTION

Genome-Wide Association Studies (GWAS) and Whole Genome Sequencing (WGS) have successfully identified common and rare variants associated with complex traits, most of which are in the noncoding genome (Hrdlickova *et al.*, 2014). However, the association between a variant and a trait does not prove the causality link because the association can result from a causal effect of the variant itself or from other variants in the same linkage disequilibrium block. To identify the disease/trait-associated causal variants is challenging due to a limited study sample size, a small effect size of variants and the linkage disequilibrium. Moreover, expression quantitative trait locus (eQTL) analysis is able to identify eQTLs in different tissues and cell types, which have a great potential to regulate targeted gene expression (Consortium, 2015). However, computationally identified eQTLs may not the true regulatory variants for gene expression. Thus, it is important to pinpoint the context-specific functional noncoding variants (NCVs) such as disease/trait-associated causal variants and tissue/cell type-specific regulatory variants.

In the past decades, thousands of functional NCVs have been discovered and deposited in database such as regulatory variants in Human Gene Mutation Database (HGMD) (Stenson *et al.*, 2020), pathogenetic variants in ClinVar (Landrum *et al.*, 2016) and somatic mutations in Catalogue of Somatic Mutations in Cancer (COSMIC) (Tate *et al.*, 2019), which provide a compendium of validated NCVs in a mixture of context. In recent years, the development of high-throughput functional assays offers an unprecedented opportunity to evaluate the functional effects of genetic variants such as massively parallel reporter assays (MPRAs) (Melnikov *et al.*, 2014) and saturating mutagenesis CRISPR-Cas9–mediated functional genomic screen (Wen *et al.*, 2020). Technically, the functional effect of a variant is measured by evaluating the molecular phenotypic change (e.g., gene expression) of the allelic alteration in different tissues and cell types. Indeed, the advance of experiment approaches leads to the discovery of validated functional NCVs in a context-specific way.

Though tremendous efforts such as HGMD, ClinVar, COSMIC and multiple wet-lab experiments have discovered thousands of functional NCVs, the scale of the discovery is still small compared to 600 million unique variants identified in gnomAD (Koch, 2020). Nevertheless, it is experimentally infeasible to evaluate all 600 millions of variants considering the technical difficulty and financial cost. Consequently, the full landscape of functional NCVs in the entire human genome is still elusive. Therefore, it is demanded to develop a computational model to *in silico* predict the functional consequence of genome-wide noncoding variants. With the rapid development of massively parallel sequencing technologies, tissue/cell typespecific epigenome, transcriptome, methylome and chromatin interaction data have been available in large national consortia such as the Encyclopedia of DNA Elements (ENCODE) (Consortium, 2004), Roadmap Epigenomics (Bernstein *et al.*, 2010) and 4D Nucleome (Dekker *et al.*, 2017). These multi-omics data provide an alternative way to define the function of a NCV by evaluating the enrichment of omics annotations or change of omics annotations under the allelic alteration.

Consequently, the integrative modeling of omics annotations and functional NCVs in HGMD, ClinVar and COSMIC has driven the development of multiple computational methods to predict genome-wide NCVs. For example, GWAVA (Ritchie *et al.*, 2014) adopts random forest, which is trained by noncoding regulatory variants in HGMD, to prioritize genome-wide NCVs; FATHMM-MKL (Rogers *et al.*, 2018) and its improved version FATHMM-XF (Shihab *et al.*, 2015) use support vector machine, which is trained by noncoding regulatory variants in HGMD with kernels designed by multi-omics feature groups, to predict both coding and noncoding pathogenic point mutations; PAFA (Zhou and Zhao, 2018) utilizes sparse logistic regression, which is trained by noncoding pathogenic variants in ClinVar and GWAS SNPs in GWASdb (Li *et al.*, 2012), to predict noncoding risk variants; ncER (Wells *et al.*, 2019) adopts XGBoost (Noh *et al.*, 2021), which is trained by noncoding regulatory variants in HGMD and noncoding pathogenetic variants in ClinVar, to predict noncoding pathogenic variants. Moreover, unsupervised methods have also been developed for their advantage of not relying on labelled functional NCVs especially when functional NCVs are few. For example, FunSeq2 (Fu *et al.*, 2014) weights integrated omics features to predict noncoding regulatory variants in cancer; GenoCanyon (Lu *et al.*, 2015) uses a mixture model to predict noncoding deleterious variants; DVAR (Yang *et al.*, 2019) uses a nonparametric Bayesian model to define omics clusters for predicting noncoding regulatory variants; fitCons (Gulko *et al.*, 2015) and its improved version FitCons2 (Gulko and Siepel, 2019) estimate the probability of fitness consequence of point mutations based on clustered functional genomic fingerprints; DeepSEA adopts a convolutional neural network-based multi-task learning framework to evaluate the allelic effects of noncoding variants by integrating multi-omics data (Zhou and Troyanskaya, 2015).

Despite the success of previous work, we are still facing many challenges to predict functional NCVs. First, the context such as disease, tissue and cell type should be considered in the prediction task because NCVs perform different biological function in different context. Second, though context-specific methods such as DIVAN (Chen *et al.*, 2016), (Chen and Qin, 2017), TIVAN (Chen *et al.*, 2019) and WEVar (Wang *et al.*, 2021) utilize surrogates of functional NCVs such as disease-specific GWAS SNPs and tissue-specific cis-eQTLs as the training set to predict noncoding disease-specific risk variants and cell-type-specific regulatory variants, these variants in the training set may not true context-specific functional NCVs due to linkage disequilibrium. Instead, it is ideal and optimal to directly use the experimentally validated functional NCVs as the training set to develop a supervised machine learning model for predicting context-specific functional NCVs. However, validated functional NCVs are rare because web-lab experiments such as MPRA and CRISPR-Cas9 are expensive, laborious, and difficult to implement. This situation imposes a challenge to adopt a classic supervised machine learning methods to use the validated functional NCVs as the training set directly, because supervised methods usually require a large training sample size to achieve a robust prediction. Thus, directly applying these methods will suffer from a small training sample size, resulting in unfavorable prediction performance.

Transfer learning is a machine learning approach, where a model developed for a source task is reused as the starting point for a model on a target task (Pan and Q., 2010). A common assumption for transfer learning is that the feature space in source task is similar to the target task. Usually, the source task has a large sample size and target task has few samples, so the target task can benefit from the source task by learning the common feature space. To date, different algorithms have been used for transfer learning with different applications in bioinformatics. For example, matrix factorization-based transfer learning discovers disease-associated patterns in rare disease by training on a large public compendia of gene expression data (Taroni *et al.*, 2019) and identifies common regulatory patterns across scATAC-seq data sets (Erbe *et al.*, 2020); Network-based transfer learning reconstructs human gene regulatory network (GRN) (Mignone *et al.*, 2020) by transferring the knowledge from mouse GRN. With the development of deep learning in recent decades, transfer learning in deep learning, which is also called deep transfer learning, have been widely applied in various domains for the flexibility and easy implementation of deep learning model. Deep transfer learning has been first used in computer vision due to the lack of sufficient labelled image data to train a new model from the scratch. The application of deep transfer learning is motivated by the nature of convolutional neural network. Specifically, the convolutional layers close to the input layer can learn low-level features such as lines and edges; middle layers can learn simple shapes such as circles, squares, ellipse; and layers close to the output layer can learn high-level features, which tend to be a unique pattern of the images for achieving image classification. Compared to high-level features, the low-level and mid-level features are more generic, which can be learnt from the source task and utilized in the target task. Adopting the idea, deep transfer learning has been widely used in the field of medical imaging and bioinformatics such as identifying novel imaging biomarkers of Alzheimer’s disease progression (Li *et al.*, 2021); imputing missing RNA-sequencing data from DNA methylation (Zhou *et al.*, 2020); predicting CYP2D6 haplotype function (McInnes *et al.*, 2020); correcting batch effects (Wang *et al.*, 2019b) and denoising single-cell RNA-seq (Wang *et al.*, 2019a).

Despite the popularity and success of transfer learning, it has been seldom applied in the field of genetics. To fill this gap, we propose TLVar, which a deep <u>T</u>ransfer <u>L</u>earning, to predict context-specific functional noncoding <u>V</u>ariants. Leveraging large-scale generic functional NCVs collected from multiple resources as the training set, we pretrain a base network, which is based on convolutional neural network. Then, we transfer the pretrained convolutional layers to a second target network with dense layers retrained using context-specific functional NCVs given the pretrained layers frozen. The trained target network aims to improve the prediction for functional NCVs from the same context. Next, we benchmark TLVar against deep learning models without pretraining or retraining, and 18 competing computational methods on three MPRA datasets and 16 GWAS datasets. In addition, we perform a comprehensive sensitivity analysis of the sample size for pretraining in the base network and retraining in the task network respectively. To our best knowledge, the work is the first time to exploit deep transfer learning in predicting functional NCVs using genomic sequence only. With the growth of MPRA/CRISPR-Cas9 validated variants in multiple tissues/cell types, TLVar will become a promising tool for accurately predicting context-specific functional NCVs, which will benefit the genetic research community.

## 2 METHODS

### 2.1 The deep transfer learning model

We adopt a convolutional neural network (Yamashita *et al.*, 2018) as the network architecture for developing deep transfer learning model “TLVar” (Fig 1). The transfer learning starts from pretraining a base network. The target network will inherit the first layers from pretrained base network with the remaining layers retrained toward the target task given pretrained layers frozen. Specifically, the base network consists of two convolutional layers and two dense layers, which is pretrained using integrated generic functional NCVs from HGMD, ClinVar and ORegAnno. In the transfer process, the two convolutional layers from base network are inherited to the target network with two dense layers retrained toward the target task using context-specific functional NCVs (e.g., cell-type specific regulatory variants, disease-specific risk variants). Moreover, we decide not to fine-tune the target network because the training sample size of context-specific functional NCVs in target network is small, which may cause overfitting. Instead, the inherited convolutional layers are frozen in the retraining process. To test, validate and train target network, we use 20% of all context-specific functional NCVs as the independent testing set, the remaining 80% NCVs as the training set among which 20% is used as the validation set. TLVar aims to improve the prediction for functional NCVs from the same context.

**Fig. 1.**
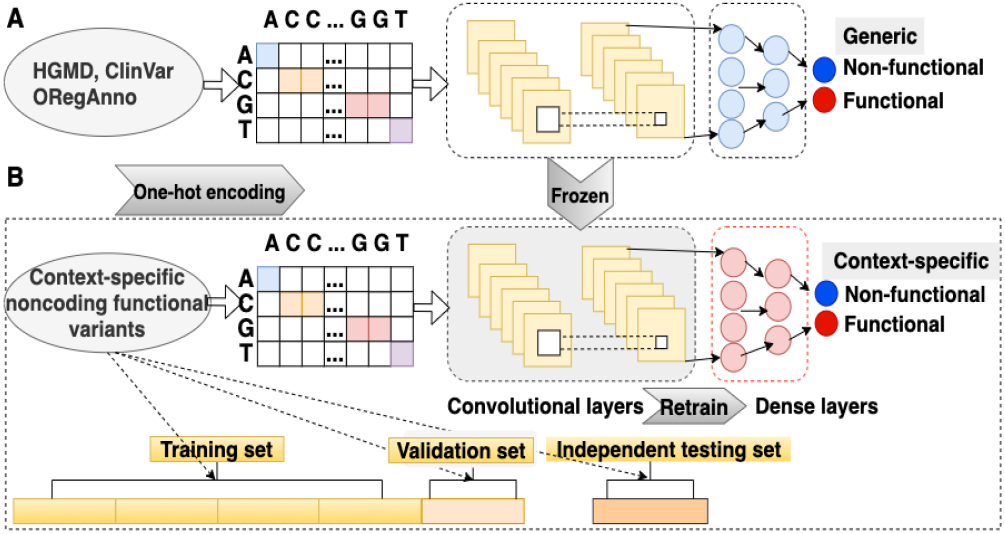
Overview for the deep transfer learning model “TLVar”. The deep transfer learning model adopts convolutional neural network, which consists of two convolutional layers and two dense layers. (A) A base network is pretrained using large-scale generic functional NCVs mainly from HGMD, ClinVar and ORegAnno and (B) transfer the pretrained convolutional layers to the target network with dense layers retrained using context-specific functional NCVs given the transferred convolutional layers frozen. To test, validate and train target network, we use 20% of all context-specific functional NCVs as the independent testing set, the remaining 80% NCVs as the training set among which 20% as the validation set. TLVar aims to improve the prediction for functional NCVs from the same context.

TLVar utilizes the flanking genomic sequence, which has the NCV as midpoint extending 500bp upstream and downstream, as the model input. The sequence is further one-hot encoding (A: [1,0,0,0], C: [0,1,0,0], G: [0,0,1,0] and T: [0,0,0,1]) as the feature representation for each NCV, which directly connects to a convolutional layer. Each convolutional layer is activated by the Rectified Linear Unit and followed by a max pooling layer. The output of last convolutional layer is flattened and connected to multiple dense layers. To improve generalization, hidden neurons in each dense layer are randomly dropped out to avoid overfitting. The final output is the binary label to indicate if the NCV is functional or not. Without loss of generality, the network architecture of the base network and target network are set the same.

### 2.2 Model training

We minimize the binary cross-entropy, which is defined as 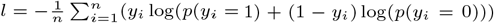, to update the network parameters using adaptive moment estimation (Adam) (DP. and J., 2014) as the optimizer in back-propagation for both pretraining the base network and retraining the target network. We train the network in 50 epochs, which demonstrate to be sufficient for model convergence. In each epoch, the whole training set is split into multiple batches and each batch pass forward and backward through the network. When the objective function converges on the validation set, we obtain the estimated network parameters. Given flanking genomic sequence of a NCV, the trained network can predict the probability of the NCV being functional.

To further increase the training efficiency, we adopt “EarlyStopping” and “ModelCheckpoint” two techniques. EarlyStopping stops the training when prediction performance on validation set stops improving, which can happen before all epochs ends. Thus, EarlyStopping can help reduce training time. To stabilize EarlyStopping, we add a delay to trigger EarlyStopping by 5 epochs when there is no improvement. Together with EarlyStopping, ModelCheckpoint saves the trained model when the prediction performance on validation set continues to improve. Therefore, the trained model with best prediction performance on the validation dataset are saved, which is not necessary the model trained in the last epoch. For hyperparameter tuning in each model, we adopt RandomSearch tuner from kerasTuner (O’Malley *et al.*, 2019). Specifically, we design a fine grid of key hyperparameters such as pooling size and kernel sizes and use RandomSearch tuner automatically to select the optimal hyperparameters.

### 2.3 Generic dataset

The generic functional NCVs for pretraining the base network are collected from an integrative collection of disease causal and regulatory variants, which are curated by Li et al. (Li *et al.*, 2016). The variant set consists of regulatory variants in HGMD (Stenson *et al.*, 2012), pathogenic variants in ClinVar (Landrum *et al.*, 2016), regulatory variants in ORegAnno (Lesurf *et al.*, 2016) and fine-mapping candidate causal SNPs for 39 diseases with a total of 5,247 positive variants and 55,923 negative variants (Farh *et al.*, 2015). These large-scale compiled variants are associated with different diseases, tissues and cell types and are in different noncoding regions including transcriptional initiation site, initiation codon, polyadenylation site or termination codon, which make them an ideal training variant set to learn low-level feature representations of NCVs.

### 2.4 MPRA datasets

We include three MPRA datasets in the analysis. In the MPRA experiment, a positive regulatory variant is defined as at least one of the two alleles of the NCV exhibits a significantly high transcriptional activity, which is measured by differential abundance of transcripts versus plasmid input. The first MPRA dataset contains 693 positive variants and 2,772 negative variants in GM12878 lymphoblastoid cells (He *et al.*, 2018). The other two MPRA datasets, which are also derived from GM12878 lymphoblastoid cells (Tewhey *et al.*, 2016), are processed by Critical Assessment of Genome Interpretation (CAGI) eQTL challenge (Kreimer *et al.*, 2017). “CAGI_train” dataset consists of 345 positive variants and 2,528 negative variants, and “CAGI_test1” dataset contains 348 positive variants and 2,460 negative variants.

### 2.5 GWAS datasets

For positive variants, we collect the noncoding GWAS SNPs (pvalue< 1 × 10^-4^) from 4 cardiovascular diseases, 9 neurological diseases and 3 immune diseases with a total number of 16 diseases in Association Results Browser (https://www.ncbi.nlm.nih.gov/projects/gapplus/sgap_plus.htm) as shown in Table S1. For each disease, using noncoding variants in 1000 Genomes (1000 Genomes Project *et al.*, 2015) as the background, we use distance to the nearest transcription starting site (TSS) as the criterion to create a set of negative variants such that distances to the nearest TSS of negative variants having a similar empirical distribution as distances to the nearest TSS of positive variants. Without loss of generality, we keep the number of negative variants the same as the number of positive variants.

### 2.6 Evaluation metrics

The prediction performance is evaluated using the area under the receiver operating characteristics (AUC) and Matthews correlation coefficient (MCC).

### 2.7 Whole-genome functional scores from 18 computational methods

To compare TLVar, we collect the genome-wide pre-computed functional scores from 18 computational methods, which include 11 supervised methods and 7 unsupervised methods (Table S2). Moreover details of these computational methods can be found in Text S1.

## 3 RESULTS

### 3.1 The deep transfer learning model

To overcome the limitation of classic supervised learning models, which usually requires a large sample size to achieve a robust prediction, we develop a supervised deep transfer learning approach “TLVar” by leveraging the powerful supervised learning and experimentally validated functional NCVs. The key hypothesis is that all functional NCVs share low-level feature representation as these variants are believed to have the key biological functions such as gene regulation. Technically, the transferred large-scale generic NCVs can stabilize the training for low-level feature representation, and few context-specific functional NCVs will learn high-level context-specific features toward the prediction in the target task. Specifically, the deep transfer learning model adopts convolutional neural network, which mainly consists of two convolutional layers and two dense layers (Fig 1). The base network is pretrained using large-scale generic functional NCVs and pretrained convolutional layers are transferred from the base network to the target network with dense layers retrained using context-specific functional NCVs given transferred convolutional layers frozen. In this way, the deep transferring model will transfer the knowledge from large-scale generic functional NCVs to improve the prediction for context-specific functional NCVs.

### 3.2 Transfer learning improves the prediction for experimentally validated regulatory variants

First, we pretrain the convolutional neural network using the generic functional NCVs from HGMD (Stenson *et al.*, 2012), ClinVar (Landrum *et al.*, 2016) and ORegAnno (Lesurf *et al.*, 2016). Next, we apply transfer learning to three MPRA datasets with a moderate size of validated noncoding regulatory variants. The first MPRA dataset (MPRA_GM12878) consists of 693 noncoding regulatory variants. The other two MPRA datasets (CAGI_train and CAGI_test1), which are collected from the CAGI eQTL challenge (Kreimer *et al.*, 2017), contain 345 and 348 noncoding regulatory variants respectively.

To demonstrate the advantage of deep transfer learning model, we compare the following deep learning models with the same network architecture, which mainly consists of two convolutional layers and two dense layers: (i) Base-model: all layers of the base network is pretrained by large-scale generic functional NCVs; (ii) Self-model: all layers of the network is trained by context-specific functional NCVs (e.g., cell type-specific regulatory variants); (iii) Deep transfer learning model (TLVar): the convolutional layers are inherited from pretrained base network and the dense layers are retrained by context-specific functional NCVs in the target network given convolutional layers frozen, where the pretrained layers are leveraged as the feature extractor. To benchmark all models, we randomly sample 20% regulatory variants as the independent testing set. To evaluate the impact of sample size of MPRA regulatory variants in training TLVar and Self-model, we construct three training sets using 10%, 50% and 100% of the remaining 80% regulatory variants. In each training set, 20% regulatory variants are randomly sampled as the validation set.

To reduce the bias from random sampling, we repeat the experiment 50 times. As a result, we report AUCs and MCCs for all methods across different levels of the training set (10%, 50% and 100%) in each MPRA dataset (Fig 2, Fig S1). In addition, we use two-sided paired Wilcoxon rank-sum test to evaluate the difference of 50 AUCs and MCCs between any two methods (Fig S2, Fig S3). Overall, Basemodel, which is trained by generic functional NCVs, demonstrates predictive power for regulatory variants. This observation indicates that functional NCVs from different context share similar low-level features. It is therefore useful to utilize large-scale generic functional NCVs to stabilize the learning for low-level features, which will improve the prediction for cell type-specific regulatory variants. Moreover, TLVar has the overall best performance. This observation indicates the advantage of using transfer learning improves the prediction for context-specific regulatory variants by leveraging both generic function NCVs.

**Fig. 2.**
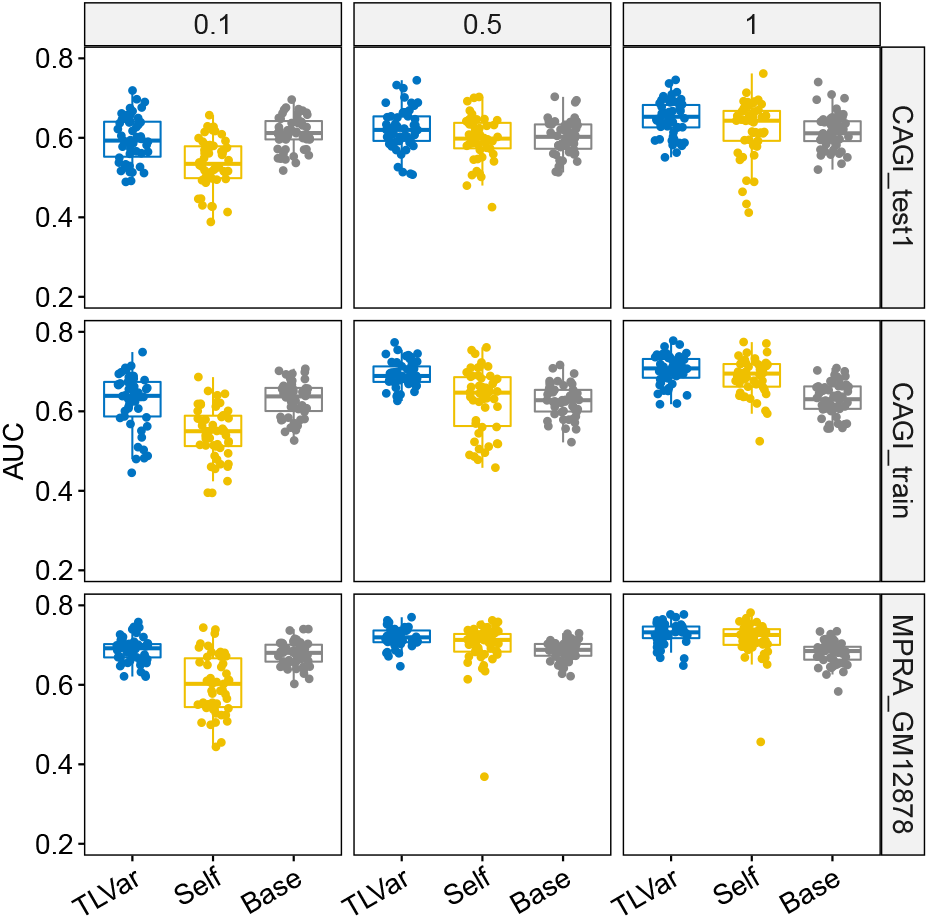
The following deep learning models are benchmarked in three MPRA datasets in terms of AUC: (i) Base-model: all layers of the base network is pretrained by large-scale generic functional NCVs; (ii) Selfmodel: all layers of the network is trained by context-specific functional NCVs; (iii) Deep transfer learning model (TLVar): the convolutional layers are inherited from pretrained base network and the dense layers are retrained by context-specific functional NCVs in the target network given convolutional layers frozen, where the pretrained layers are leveraged as the feature extractor. To benchmark all models, we randomly sample 20% MPRA regulatory variants as the independent testing set. To evaluate the impact of sample size of MPRA regulatory variants, we construct three training sets using 10%, 50% and 100% of the remaining 80% regulatory variants. In each training set, 20% regulatory variants are randomly sampled as the validation set.

When the full training set (100%) is used for retraining dense layers, TLVar has the best performance in terms of medians of AUCs followed by Self-model and Base-model in three datasets (CAGI_test1: 0.653, 0.643, 0.612; CAGI_train: 0.708, 0.696, 0.630; MPRA_GM12878: 0.732, 0.726, 0.685). The improvement of TLVar is also significant, which is demonstrated by pairwise comparison between TLVar and any other method (Fig S2). The advantage of TLVar to Base-model proves that the dense layers trained by the regulatory variants will improve the prediction than trained by generic functional NCVs because the regulatory variants can learn high-level features specifically for the target prediction task. Moreover, the improvement of TLVar to Self-model validates that using large-scale generic functional NCVs will improve the learning for low-level features than using few regulatory variants. In other words, lacking sufficient sample size of regulatory variants can deteriorate the prediction even if the training and testing variants are from the same context because a large sample size is required for model convergence and learning network parameters correctly for convolutional layers. Furthermore, Self-model significantly outperforms Base-model in terms of AUCs, which may benefit from the context-matching between training variants and testing variants (Fig S2). Overall, these observations indicate that transfer learning boosts the predictive power by leveraging the strength of large-scale generic functional NCVs pretraining convolutional layers to learn the shared low-level features and regulatory variants retraining dense layers to learn high-level context-specific features if the training set of regulatory variants is reasonably large.

Next, when the training set of regulatory variants for dense layers decreases, we find that the performance of Base-model remains stable. The rationale is that Base-model is not affected by the sample size of regulatory variants because the dense layers of Base-model are trained using generic functional NCVs. As expected, the performance of TLVar and Self-model declines because the sample size affects the training for dense layers of TLVar and all layers of Self-model. As the sample size decreases, the model tends to be underfitting and the prediction performance is deteriorated. For TLVar, the median AUC decreases from 0.653 (100%) to 0.620 (50%) to 0.593 (10%) in CAGI_test1 dataset; from 0.708 (100%) to 0.689 (50%) to 0.640 (10%) in CAGI_train dataset; from 0.732 (100%) to 0.720 (50%) to 0.692 (10%) in MPRA_GM12878 dataset. For Self-model, the median AUC decreases from 0.643 (100%) to 0.598 (50%) to 0.535 (10%) in CAGI_test1 dataset; from 0.700 (100%) to 0.647 (50%) to 0.550 (10%) in CAGI_train dataset; from 0.726 (100%) to 0.713 (50%) to 0.603 (10%) in MPRA_GM12878 dataset. For TLVar, the adjacent differences of AUCs between different training size are 0.033 (100% vs 50%), 0.027 (50% vs 10%) in CAGI_test1 dataset; 0.019 (100% vs 50%), 0.059 (50% vs 10%) in CAGI_train dataset; 0.012 (100% vs 50%), 0.028 (50% vs 10%) in MPRA_GM12878 dataset. For Self-model, the adjacent differences of AUCs between different training size are 0.045 (100% vs 50%), 0.063 (50% vs 10%) in CAGI_test1 dataset; 0.053 (100% vs 50%), 0.097 (50% vs 10%) in CAGI_train dataset; 0.013 (100% vs 50%), 0.11(50% vs 10%) in MPRA_GM12878 dataset. Comparing the declining performance between TLVar and Self-model, we find that TLVar is less sensitive than Self-model to the decreasing sample size. Especially, the performance of Self-model declines more dramatically when the training set decreases from 50% to 10% compared to TLVar. These observations are further confirmed by statistical significance comparison between TLVar and Self-model, where the advantage of TLVar over Self-model measured by pvalue grows more significantly when the training size decreases (Fig S2). Moreover, the AUC distribution of TLVar is less dispersed compared to Self-model with the decrease of sample size especially for CAGI_train and MPRA_GM12878 datasets. Overall, these observations indicate that TLVar is more robust than Self-model when the training sample size decreases. The robustness of TLVar can be explained by only dense layers need to be trained for TLVar but both convolutional layers and dense layers need to be trained for Self-model. Thus, diminishing training sample size will cause more underfitting of Self-model than TLVar, which results in more severely deteriorated performance.

Regarding the performance comparison among other methods, TLVar loses the advantage to Base-model when sample size diminishes to 10% in CAGI_test1 dataset (0.593 vs 0.612). Self-model is outperformed by Base-model in CAGI_test1 dataset (0.598 vs 0.603) when the training size decreases to 50%, and in all datasets when the training size decreases to 10% (CAGI_test1: 0.535 vs 0.612; CAGI_train: 0.550 vs 0.638; MPRA_GM12878: 0.603 vs 0.681). The reason may be because the benefit from the context-matching between training variants of dense layers and testing variants is offset by the underfitting caused by the decreased sample size of training variants. Besides AUC, we also examine the performance measured by MCC (Fig S1, S3). The overall trend is similar to AUC. Therefore, we conclude that transfer learning indeed improves the prediction for regulatory variants based on the evaluation of AUC and MCC across different training size in three MPRA datasets.

### 3.3 Transfer learning improves the prediction for disease-associated risk variants

To further evaluate the performance of TLVar to predict context-specific functional NCVs, we apply TLVar to predict noncoding disease-specific risk variants in 16 GWAS datasets across three disease classes, which include neurological diseases, cardiovascular diseases and immune diseases. The sample size of disease-specific risk SNPs ranges from 80 to 725 with a median 258 (Table S1), which is ideal to test the performance of TLVar. To construct the negative SNPs, we restrict the negative SNPs to have a similar empirical distribution of distance to nearest TSS as the GWAS SNPs, which will balance the regulatory potential between positive and negative set.

The deep learning model architecture, the training and testing procedure are set the same as aforementioned. Specifically, 20% of whole dataset is used as independent testing, 80% is used as training set among which 20% is used as validation set. To control the bias from random sampling, the experiment is repeated 50 times. For each dataset, we report the AUCs and MCCs from 50 experiments for each method when the training set varies from 10%, 50% and 100%, and perform method comparison using two-sided paired Wilcoxon rank-sum test (Fig S4-S9). We also report the median of AUCs and MCCs from 50 experiments in each dataset for each method across different levels of training set (Fig 3). Since the overall trends for AUCs and MCCs for all methods across all datasets are similar, we focus on using AUCs for model comparison.

**Fig. 3.**
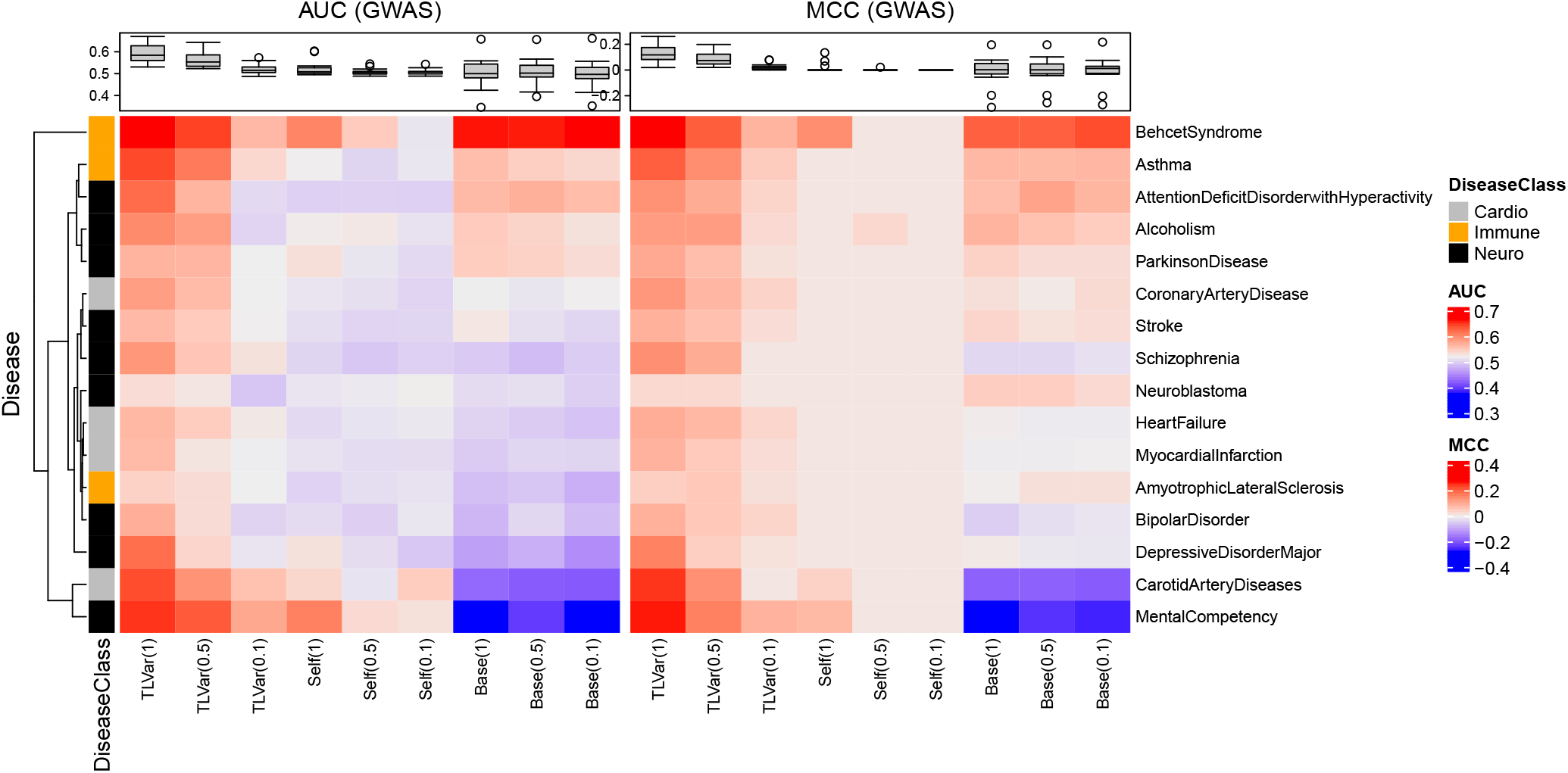
Three deep learning models (TLVar, Self-model, Base-model) on different levels of training set (10%, 50% and 100%) for retraining dense layers are compared to predict disease-specific risk variants in 16 GWAS datasets across three disease classes (neurological diseases, cardiovascular diseases and immune diseases) in terms of AUC and MCC.

First, we perform a global evaluation to compare any two methods using twosided paired Wilcoxon rank-sum test on the median AUCs of across 16 diseases in 50 experiments. As expected, the AUCs for TLVar declines when training set decreases. Overall, TLVar has the best performance, followed by Self-model and Base-model. When using 100% of the training set, TLVar has a clear advantage over Self-model and Base-model. (pvalue=4.814 × 10^-4^ for TLVar vs Self-model; pvalue=3.052 × 10^-5^ for TLVar vs Base-model). This advantage still holds when the training set decreases to 50%. However, when the training set decreases to 10%, TLVar only significantly outperforms Self-model (pvalue=0.023 for TLVar vs Self-model, pvalue=0.256 for TLVar vs Base-model).

Next, we examine the method comparison stratified by the disease class. Overall, TLVar maintains good performance in Behcet’s Syndrome, Asthma, Mental Competency and Carotid Artery Disease (Fig 3). In contrast, Base-model performs well in Behcet’s Syndrome but performs poorly in Mental Competency and Carotid Artery Disease (Fig 3). For neurological diseases (Fig S4), TLVar has consistently better performance than all methods when the training set is 100%. When the training set declines to 50%, TLVar still holds a clear advantage in Alcoholism, Bipolar Disorder, Mental Competency, Parkinson’s Disease, Schizophrenia and Stroke. For cardiovascular diseases (Fig S6), TLVar outperforms Self-model and Base-model in all diseases when the training set is 100% and the trend still holds when the training set drops to 50%. For immune diseases (Fig S8), when the training set is 100% and 50%, TLVar holds an advantage over both Self-model and Base-model except for Basemodel in Behcet’s syndrome (50%). However, when the training set decreases to 10%, the advantage of TLVar over other methods diminishes significantly in all diseases.

Overall, TLVar performs better than other methods to predict disease-specific risk variants especially when the training size is reasonable large (50%, 100%). When the training set decreases to 10%, TLVar may suffer from underfitting, resulting in deteriorated performance.

### 3.4 Transfer learning outperforms 18 machine learning methods in predicting experimentally validated regulatory variants

We’ve demonstrated that TLVar improves the prediction for noncoding regulatory variants compared to deep learning models without using transfer learning. However, TLVar only utilize the variant-centered flanking genomic sequence as input features. Besides genomic sequence, existing computational methods also adopt multi-omics functional data such as epigenomic, transcriptomic and chromatin interaction data as input features. To compare TLVar to the competing approaches using either genomic sequence or multi-omics functional data as input features, we collect functional scores from 18 computational methods, which include 11 supervised methods and 7 unsupervised methods. These functional scores measure the regulatory or deleterious potential of NCVs (Table S2).

We adopt a five-fold cross-validation approach for evaluating all methods. Specifically, the whole training set is split into five-folds. Each one-fold is used iteratively as independent testing set, and the remaining four-folds is used as the training set among which 20% variants are randomly sampled as the validation set. Thus, prediction scores for the one-fold can be obtained by the trained model. Similarly, precomputed scores for the one-fold can be obtained from other methods directly. For each method, the scores are merged across five-folds, and compared to the true labels of regulatory variants for calculating AUCs and MCCs.

As a result (Fig 4), TLVar tops other methods in terms of AUC in CAGI_train and MPRA_GM12878 two datasets, and on par with GenoCanyon in CAGI_test1 dataset. Especially, TLVar achieves a significant improvement of AUC in CAGI_train dataset. Similarly, TLVar significantly outperforms other methods in terms of MCC in CAGI_train and MPRA_GM12878 two datasets, and slightly secondary to GenoCanyon in CAGI_test1 dataset. Three unsupervised methods, which include FunSeq2, DVAR and GenoCanyon, are consistently among the top in all datasets. The superior performance of the three methods can be explained by the benefit from omics integration and the nature of unsupervised learning, which is less sensitive to training labels. GenoCanyon integrates genomic conservation score, open chromatin, histone modification and transcription factor binding to identify regulatory potential of NCVs. Similar to GenoCanyon, DVAR utilizes omics features to develop a functional score for NCVs. Though FunSeq2 is originally designed to prioritize noncoding regulatory variants in cancer, the integrated omics features such as ultra-conserved element, GERP scores and motif-breaking/gaining score, have a general purpose to identify functional NCVs. In contrast, CScape and CDTS consistently perform poorly across all datasets. The poor performance of CDTS may be explained by using genomic sequence only without considering functional omics data, and the deteriorated performance of CScape may be attributed to a lack of context-matching training set. CScape uses cancer mutations in COSMIC as training set. However, being cancerous is not necessarily being regulatory. Interestingly, DANN and DeepSEA, which are two deep learning methods, has moderate performance in all datasets. Overall, these observations demonstrate that using genomic sequence only and leveraging context-matching regulatory variants, TLVar holds a clear advantage than competing methods, which are developed either using genomic sequence or multi-omics functional data, to predict noncoding regulatory variants.

**Fig. 4.**
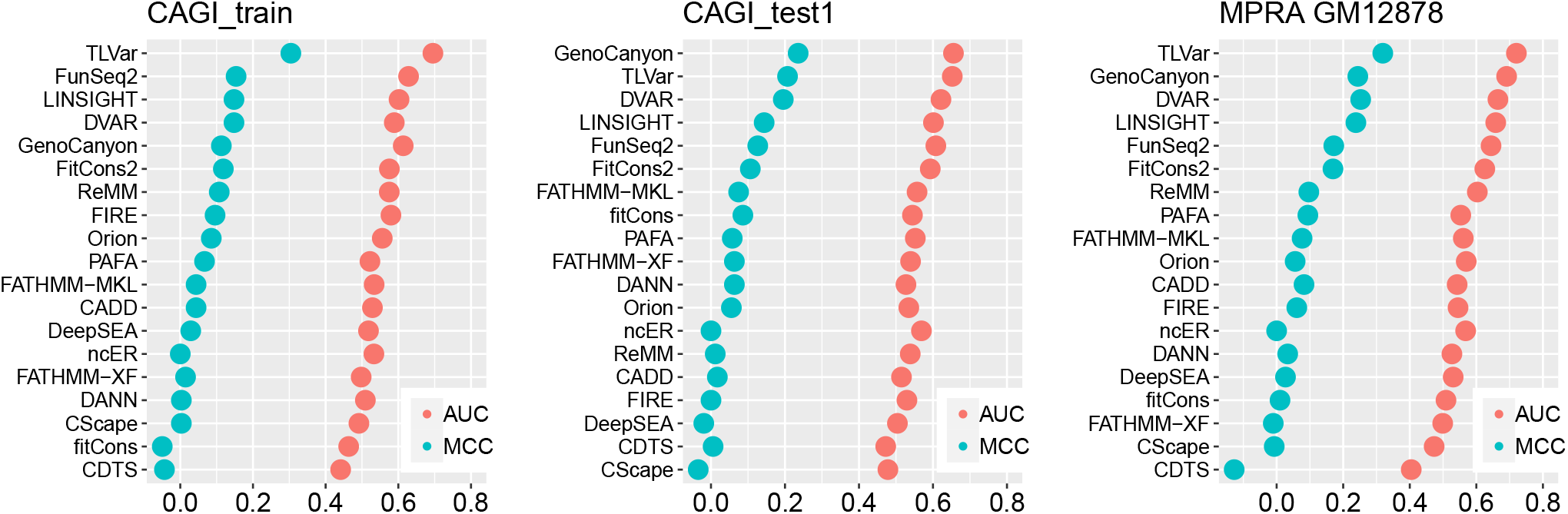
Deep transfer learning model TLVar is compared to 18 computational methods, which is based on precomputed functional scores, to predict regulatory variants in three MPRA datasets in terms of AUC and MCC using five-fold cross validation.

### 3.5 Transfer learning outperforms 18 machine learning methods in predicting disease-associated risk variants

We adopt the same aforementioned five-fold cross-validation strategy for all methods to predict disease-associated risk variants. First, we calculate AUC and MCC for each method in each GWAS dataset (Fig 5). Next, we perform two-sided paired Wilcoxon rank-sum test to evaluate the difference of AUCs and MCCs across 16 GWAS datasets between TLVar and any other method.

**Fig. 5.**
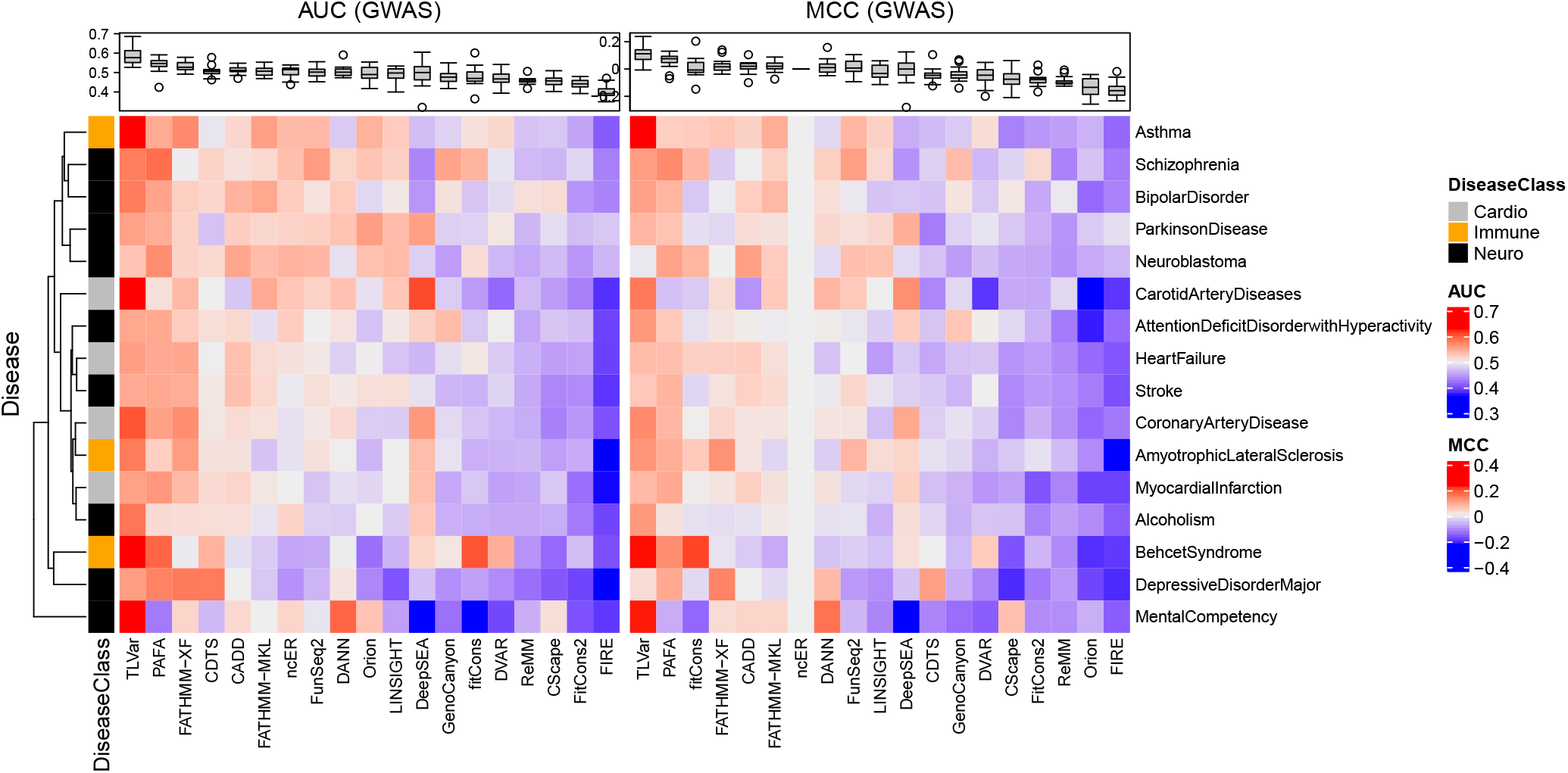
Deep transfer learning model TLVar is compared to 18 computational methods, which are based on precomputed functional scores, to predict diseasespecific risk variants in 16 GWAS datasets across three disease classes (neurological diseases, cardiovascular diseases and immune diseases) in terms of AUC and MCC using five-fold cross validation.

As a result, TLVar consistently outperforms other methods in terms of both AUC and MCC. Following TLVar, PAFA and FATHMM-XF have second best performance. This observation indicates that supervised models trained by noncoding pathogenetic variants in HGMD and/or ClinVar can be powerful to predict noncoding GWAS SNPs since both sets of variants are context-matching of being disease-associated. Moreover, FIRE, ReMM, CScape, fitCons, FitCons2, DVAR, GenoCanyon have overall poor performance. This is because FIRE is trained using highly confident cis-QTLs and thus aim to predict regulatory variants. However, regulatory variants are not necessarily risk variants. ReMM is limited to predict pathogenic regulatory variants in Mendelian diseases, therefore may be less powerful to predict risk variants associated with complex diseases. Unsupervised methods, which include fitCons, FitCons2, DVAR and GenoCanyon, are less powerful for predicting noncoding risk variants. In addition, DANN and DeepSEA, which are two deep learning methods, has moderate performance.

Comparing the prediction performance for MPRA regulatory variants and disease-associated risk variants, we find that top-performed methods in MPRA datasets including FunSeq2, DVAR and GenoCanyon have a significant performance decline in GWAS datasets. FunSeq2 has moderate performance, and DVAR, GenoCanyon have relatively poor performance in GWAS datasets. Additionally, CScape, which is a bottom-performed method in MPRA datasets, also has poor performance in GWAS datasets. These observations demonstrate that methods developed for predicting noncoding regulatory variants may not be suitable for predicting noncoding deleterious variants, and vice versa. Therefore, it is important to customize computational models to optimize the prediction for context-specific functional NCVs. Indeed, TLVar holds the advantage to be customized by functional NCVs from different context in dense layers with a limited demand of training set to achieve optimal performance.

### 3.6 Evaluation of pretraining on the effectiveness of deep transfer learning

We have demonstrated that the sample size of context-specific functional NCVs for retraining dense layers affects the prediction performance of TLVar, that is, the larger sample size of context-specific functional NCVs, the better prediction performance. Here, we further explore whether the sample size of generic functional NCVs for pretraining convolutional layers will have an impact on the prediction performance.

Without loss of generality, we perform the model evaluations on the three MPRA datasets as GWAS datasets can be evaluated in a similar way. As before, we randomly sample 20% of regulatory variants as the independent testing set and remaining 80% as the training set among which 20% is used as the validation set for retraining dense layers in TLVar and all layers in Self-model. To control the sampling bias, all experiments are repeated 50 times. In addition to TLVar, Self-model and Base-model, we include TLVar00, which has untrained convolutional layers with randomly initialized weights but retrained dense layers given convolutional layers frozen. As a result, we report the median of AUCs in all experiments, and compare all deep learning methods using twosided paired Wilcoxon rank-sum test to evaluate the relative contribution of pretrained convolutional layers and retrained dense layers (Fig 6).

**Fig. 6.**
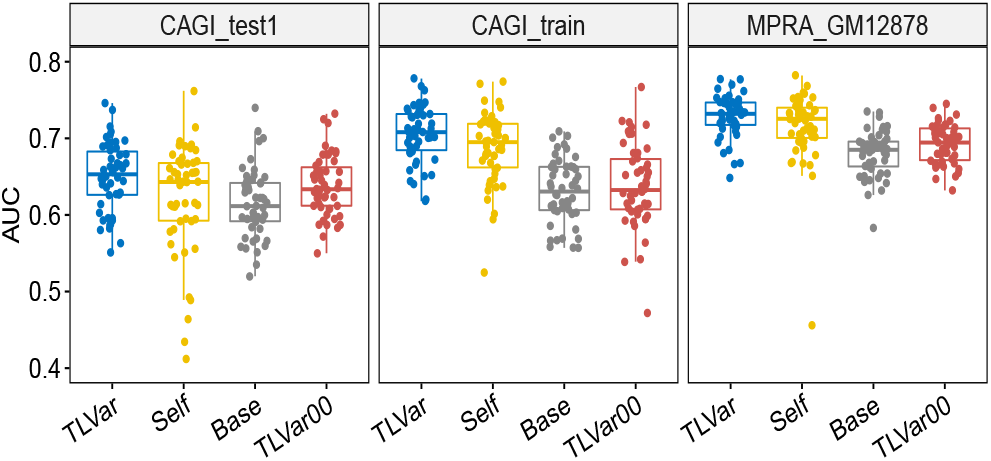
A new deep learning method TLVar00, which is the same as TLVar except the convolutional layers are not pretrained, is compared to aforementioned deep learning models in terms of AUC. We randomly sample 20% regulatory variants as the independent testing set for all models and remaining 80% as the training set among which 20% is used as the validation set. To control the sampling bias, all experiments are repeated 50 times.

As expected, TLVar significantly outperforms TLVar00 in CAGI_train and MPRA_GM12878 datasets (0.708 vs 0.633 (pvalue=7.781 × 10^-8^), 0.732 vs 0.695 (pvalue=6.157 × 10^-7^) and slightly performs better than TLVar00 in CAGI_test1 dataset (0.653 vs 0.634) (Fig S10). The observation demonstrates that pretraining convolutional layers will improve the prediction performance. Moreover, Self-model holds a clear advantage to TLVar00 in CAGI_train and MPRA_GM12878 datasets (0.696 vs 0.633 (pvalue=8.332 × 10^-6^), 0.726 vs 0.695 (pvalue=2.997 × 10^-4^)) and slightly outperforms TLVar00 in CAGI_test1 dataset (0.643 vs 0.634). This observation can be explained by the following reason. On one hand, there may be more parameters to be trained for Self-model than TLVar00. On the other hand, the untrained convolutional layers of TLVar00 causes noise in model. Therefore, the performance comparison between Self-model and TLVar00 is a comparison between the potential underfitting caused by more parameters in Self-model and potential underfitting caused by untrained convolutional layers in TLVar00. Clearly, the level of underfitting is more severe in TLVar00 than Self-model, which results in a more unfavorable performance of TLVar00. Nevertheless, TLVar00 achieves a significantly higher AUC than Base-model in CAGI_test1 and MPRA_GM12878 datasets (0.634 vs 0.612 (pvalue=0.012), 0.695 vs 0.685 (pvalue=0.048)) but has a comparable performance to Base-model in CAGI_train dataset (0.633 vs 0.631). This observation indicates the power of using context-specific functional NCVs to retrain dense layers if the testing NCVs are from the same text, even if the convolutional layers are left untrained.

## 4 DISCUSSION

Motivated by the existing gap of lacking sufficient experimentally validated functional variants to train a robust supervised learning model to predict genome-wide functional noncoding variants (NCVs), we develop a deep transfer learning model “TLVar”, which adopts convolutional neural network, to predict context-specific functional NCVs (e.g., cell-type specific regulatory variants, disease-specific risk variants). TLVar utilizes one-hot encoding variant-centered flanking genomic sequence as the feature representation of NCVs, pretrains convolutional layers utilizing a large-scale generic functional NCVs from multiple resources and retrains dense layers using small-scale context-specific functional NCVs. In this way, TLVar leverages both generic functional NCVs to learn common low-level features and context-specific functional NCVs to learn high-level features for improving the prediction of functional NCVs in the same context.

We benchmark TLVar against three deep learning models with the same network architecture, which include model with all layers trained by large-scale generic functional NCVs (Base-model); and model with all layers trained by context-specific functional NCVs (Self-model) on three MPRA datasets and 16 GWAS datasets spanning three disease class, which include neurological diseases, cardiovascular diseases and immune diseases. As a result, TLVar has an overall best performance in terms of AUC and MCC. Self-model has the secondary best performance followed by Base-model. The superiority of TLVar to Self-model demonstrates that large-scale generic functional NCVs improves the learning for network parameters in convolutional layers than few context-specific functional NCVs. The advantage of TLVar to Basemodel indicates that the dense layers retrained by context-specific functional NCVs will improve the prediction because context-specific functional NCVs can learn high-level features specifically for the target prediction task.

We also evaluate two factors, which may influence the prediction performance of TLVar. One factor is the sample size of context-specific functional NCVs for retraining the dense layers. Consequently, we find that the performance of TLVar is deteriorated with the decrease of sample size for retraining dense layers. Compared to Self-model, TLVar is more robust to the decrease of sample size for retraining dense layers. This is because given the same context-specific functional NCVs, more parameters are needed to train both convolutional layers and dense layers in Self-model than train only dense layers in TLVar given convolutional layers are frozen. In other words, using large-scale generic functional NCVs will improve the learning for low-level features in convolutional layers than using few context-specific functional NCVs. Thus, diminishing training sample size of context-specific functional NCVs will cause more underfitting in Self-model than TLVar, which results in more deterioration of prediction performance. The other factor is the sample size of generic functional NCVs for pretraining the convolutional layers. Similarly, we observe that model without pretrained convolutional layers (TLVar00) will have deteriorated prediction performance. Therefore, all the experiments validate that TLVar benefits from both pretraining convolutional layers to learn low-level features and retraining dense layers to learn context-specific high-level features for achieving an improved performance.

Nowadays, multiple computational methods, either supervised or unsupervised, have been developed for predicting functional NCVs. These approaches aim to identify noncoding regulatory/deleterious variants by utilizing genomic sequence and multi-omics annotations. Consequently, we collect the precomputed functional scores from 18 competing methods and benchmark TLVar against them in three MPRA datasets and 16 GWAS datasets. As a result, TLVar outperforms these methods in most cases, which demonstrates the advantage of using deep transfer learning to predict context-specific functional NCVs, even if the transfer learning only utilizes the variant-located flanking genomic sequence as the feature representation.

We can further improve the deep transfer learning model by enhancing the feature representation and optimizing the design of transferred layers. In this work, we use one-hot encoding genomic sequence as feature representation of NCVs for the model input. However, previous work (Chen *et al.*, 2016, 2019) demonstrates that multi-omics annotations will also improve the prediction for functional NCVs. Here, we use convolutional neural network as the algorithm for transfer learning model. To fully take the advantage of convolutional neural network, which is a flexible architecture to accommodate multi-modal input, we plan to integrate both genomic sequence and multi-omics functional annotations as the feature representations for NCVs. The multi-omics annotations can include but not limited to conservation scores, tissue-/cell type-specific epigenomic, transcriptomic and chromatin interaction features. Specifically, we plan to extract sequence features from one-hot encoding flanking sequence using convolutional layers, extract omics features from multi-omics annotations using embedding layers, and integrate both feature representations using concatenate layers before making the final prediction. Moreover, in this work, we use convolutional layers for pretraining and dense layers for retraining. However, this design may not be optimal as pretrained layers can include dense layers and retrained layers can contain convolutional layers. In the future work, we will optimize the design of pretrained layers and retrained layers by iterating through all combinations of convolutional layers and dense layers. The optimal design can be decided based on cross validation.

In this work, the deep transfer learning aims to improve the prediction for context-specific functional NCVs such as MPRA validated regulatory variants and disease-specific risk variants by transferring the knowledge from large-scale generic functional NCVs. Besides this application, the deep transfer learning model can be extended for cross-trait noncoding causal variant prediction. Particularly, for genetically correlated traits, noncoding causal variants from correlated secondary traits can be used to develop a base network to improve the prediction for noncoding causal variants from the primary trait. For example, noncoding causal variants from Autism Spectrum Disorder, Bipolar Disorder and Schizophrenia can be used as secondary traits to develop a base network to predict noncoding causal variants from Alzheimer’s Disease. We will focus on this direction in the future work.

## FUNDING

This work was supported by Indiana University Precision Health Initiative, Showalter Research Trust Fund and National Institute of General Medical Sciences of the National Institutes of Health under Award Number R35GM142701 to LC.

## Notes

### Competing Interest Statement

The authors have declared no competing interest.

