## SupplementalFile for "Exploiting deep transfer learning for the prediction of functional noncoding variants using genomic sequence"

#### 1 Supplementary tables

Table S1: Summary of 16 GWAS datasets collected from Association Results Browser

| **Disease class** | **Disease** | **#SNPs** |
|---|---|---|
| Cardiovascular | Carotid Artery Diseases | 80 |
| Cardiovascular | Coronary Artery Disease | 585 |
| Cardiovascular | Heart Failure | 530 |
| Cardiovascular | Myocardial Infarction | 584 |
| Immune | Amyotrophic Lateral Sclerosis (ALS) | 197 |
| Immune | Asthma | 252 |
| Immune | Behcet's Syndrome | 229 |
| Neurology | Alcoholism | 264 |
| Neurology | Attention Deficit Disorder with Hyperactivity (ADHD) | 199 |
| Neurology | Bipolar Disorder | 274 |
| Neurology | Depressive Disorder Major | 86 |
| Neurology | Mental Competency | 114 |
| Neurology | Neuroblastoma | 268 |
| Neurology | Parkinson Disease | 324 |
| Neurology | Schizophrenia | 238 |
| Neurology | Stroke | 725 |

Table S2: Summary of machine learning methods for providing whole-genome functional scores

| **Supervised** | **Model** | **Training set** |
|---|---|---|
| FATHMM-MKL | SVM | pathogenetic variants in HGMD |
| FATHMM-XF | SVM | pathogenetic variants in HGMD |
| CADD | SVM | high-frequency human-derived alleles |
| DANN | DNN | high-frequency human-derived alleles |
| LINSIGHT | GLM | SNPS in 69 Genomes Data (http://www.completegenomics.com/public-data/69-Genomes/) |
| FIRE | random forest | highly confident cis-eQTL SNVs |
| ncER | XGBoost | pathogenetic variants in ClinVar and HGMD |
| PAFA | sparse logistic regression | pathogenic variants in ClinVar and GWAS SNPs in GWASdb[10] |
| CScape | Classification models in PyML[1] and scikit-learn[12] | pathogenic variants in COSMIC |
| ReMM | random forest | curated pathogenic regulatory variants in Mendelian disease |
| DeepSEA | CNN | ENCODE and Roadmap Epigenomics data |
| **Unsupervised** | **Model** | |
| fitCons | Evolutionary model | |
| FitCons2 | Evolutionary model | |
| DVAR | Nonparametric Bayesian model | |
| FunSeq2 | Weighted scoring | |
| CDTS | Observed variation from expected variation | |
| Orion | Observed site frequency spectrum from the expected | |
| GenoCanyon | Mixture model | |

### 2 Supplementary text

In Table S2, for supervised methods, FATHMM-MKL [15] adopts support vector machine, which is trained by pathogenetic variants in HGMD to predict both coding and noncoding pathogenic point mutations. FATHMM-XF [16] is an extended version of FATHMM-MKL by including more training features. CADD [9] uses support vector machine, which is trained to differentiate high-frequency human-derived alleles from simulated variants to predict both coding and noncoding pathogenic variants. DANN [13] is an extension of CADD by using deep neural network to better capture non-linear relationships among features. LINSIGHT [7] utilizes generalized linear model to predict deleterious NCVs by deriving a fitness consequence score. FIRE [8] adopts random forest, which is trained by highly confident *cis*-eQTL SNVs to predict both coding and noncoding regulatory variants. ncER [18] uses XGBoost, which is trained by noncoding pathogenic variants in ClinVar and HGMD to predict noncoding pathogenic variants. PAFA [21] adopts sparse logistic regression, which is trained by pathogenic variants in ClinVar and GWAS SNPs in GWASdb to predict noncoding risk variants. CScape [14] is trained using pathogenic variants in COSMIC to predict coding and noncoding somatic point mutations. ReMM [17] adopts random forest to predict pathogenic regulatory variants in Mendelian disease. DeepSEA adopts a convolutional neural network-based multi-task learning framework to evaluate the allelic effects of noncoding variants by integrating multi-omics data [20].

For unsupervised methods, GenoCanyon [11] adopts the mixture model to predict noncoding deleterious variants. fitCons [4] estimates the probability of fitness consequence of variants via clustered functional genomic fingerprints. FitCons2 [5] is an improved version of fitCons by considering both clustered functional genomic fingerprints and evolutionary properties. DVAR [19] utilizes nonparametric Bayesian model to define clusters among omics features to predict noncoding regulatory variants. FunSeq2 [3] calculates a weighted score of integrated omics features to predict noncoding regulatory variants in cancer. CDTS [2] calculates the observed variation from expected variation to define the functional predictive score for human noncoding genome. Orion [6] calculates the observed site frequency spectrum from the expected to define the functional score for human noncoding genome intolerance to variations.

### 3 Supplementary figures

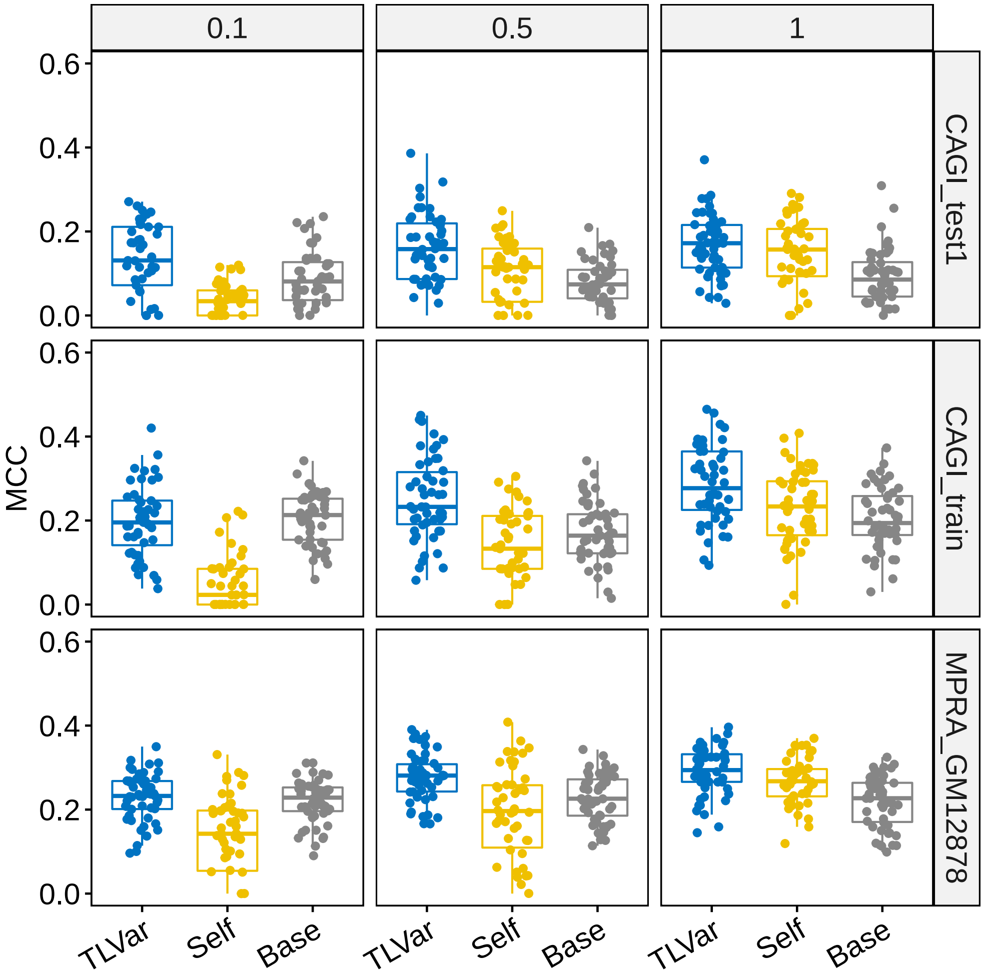


Figure S1: The following deep learning models are compared to predict context-specific MPRA variants in three MPRA datasets in terms of MCC: (i) Base-model: all layers of the base network is pretrained by large-scale generictwo-sided functional NCVs; (ii) Self-model: all layers of the network is trained by context- specific functional NCVs; (iii) Deep transfer learning model (TLVar): the convolutional layers are inherited from pretrained base network and the dense layers are retrained by context-specific functional NCVs in the target network given convolutional layers frozen, where the pretrained layers are leveraged as the feature extractor. To benchmark all models, we randomly sample 20% MPRA regulatory variants as the independent testing set. To evaluate the impact of sample size of MPRA regulatory variants in training TLVar and Self-model, we construct three training sets using 10%, 50% and 100% of the remaining 80% regulatory variants. In each training set, 20% regulatory variants are randomly sampled as the validation set.

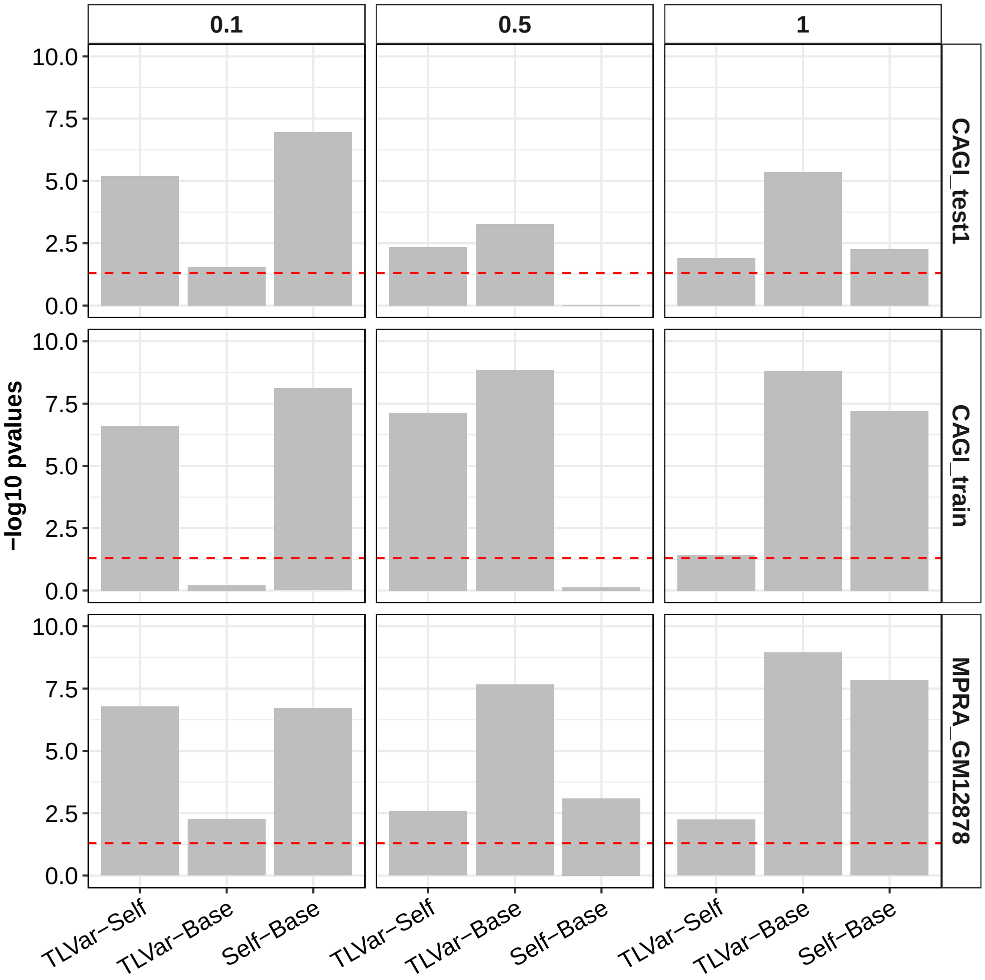


Figure S2: For three MPRA datasets, two-sided paired Wilcoxon rank-sum test is used to test the difference of AUCs in 50 experiments between two deep learning methods among TLVar, Base-model and Self-model. The red dash line is the threshold of $-log_{10}(0.05)$.

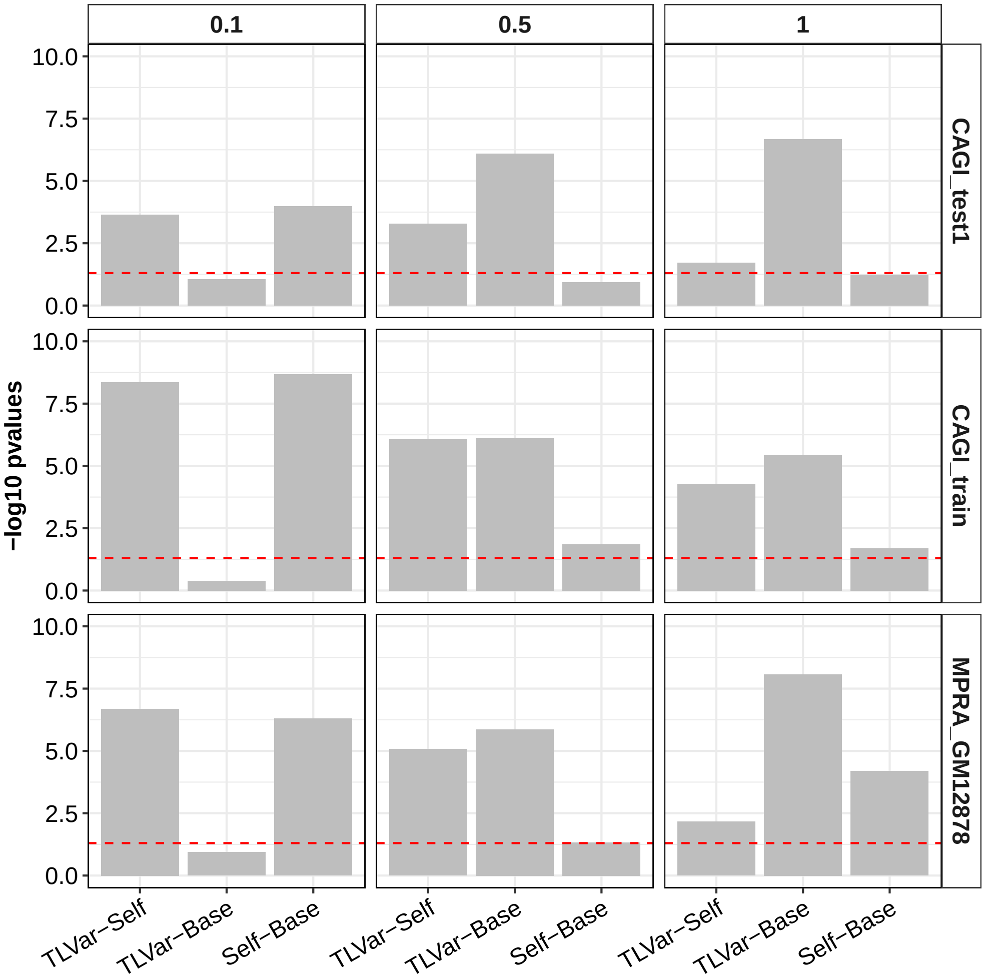


Figure S3: For three MPRA datasets, two-sided paired Wilcoxon rank-sum test is used to test the difference of MCCs in 50 experiments between two deep learning methods among TLVar, Base-model and Self-model. The red dash line is the threshold of $-log_{10}(0.05)$.

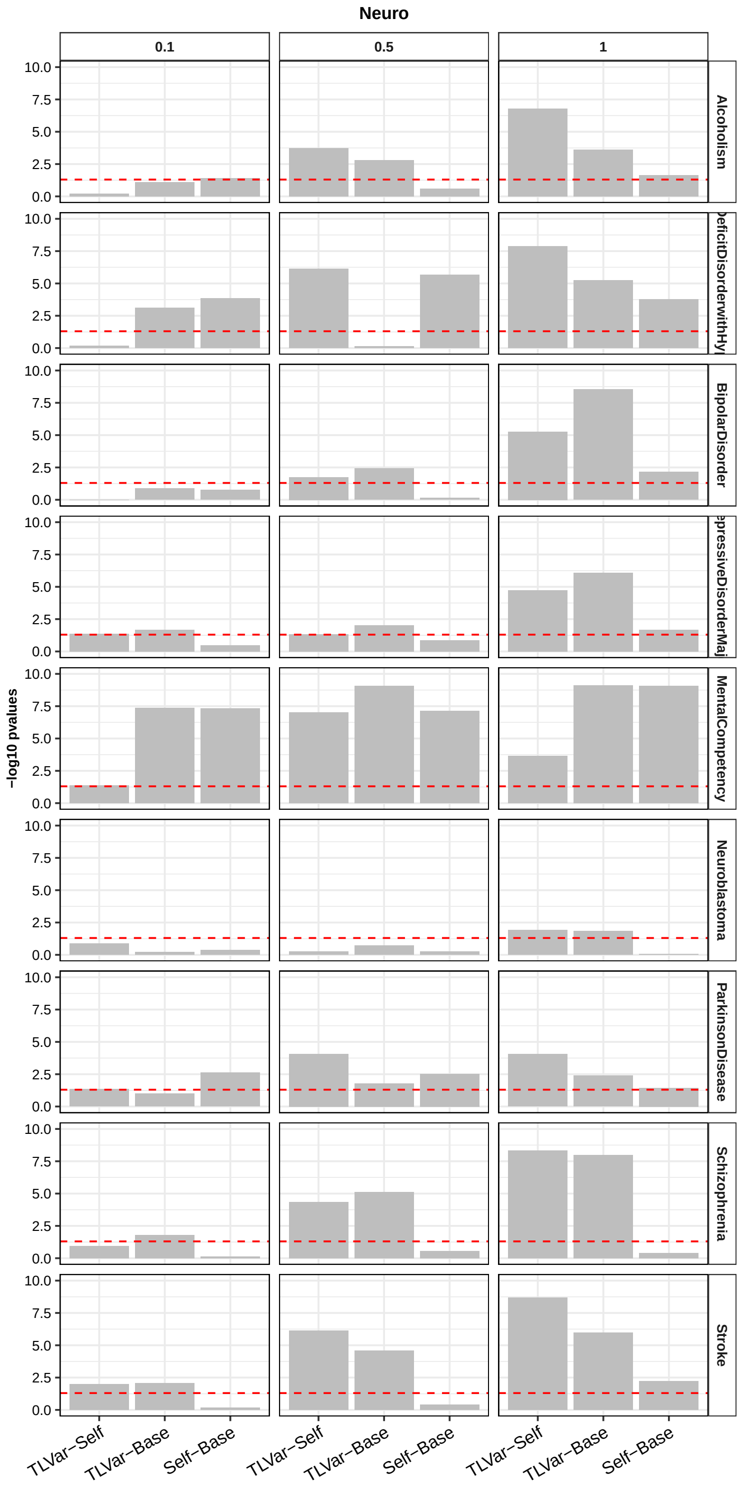


Figure S4: For 9 GWAS dataset in neurological diseases, two-sided paired Wilcoxon rank-sum test is used to test the difference of AUCs in 50 experiments between two deep learning methods among TLVar, Base-model and Self-model. The red dash line is the threshold of $-log_{10}(0.05)$.

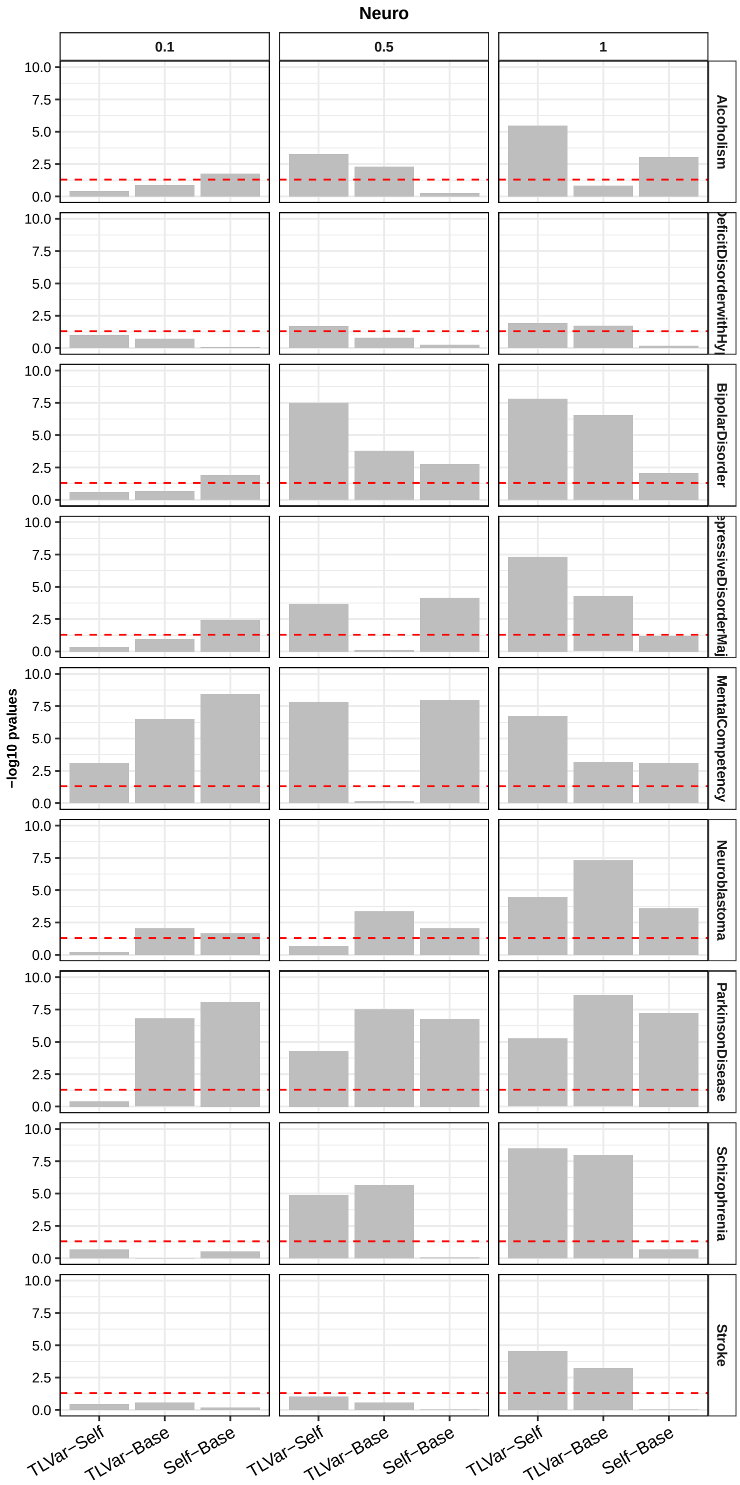


Figure S5: For 9 GWAS dataset in neurological diseases, two-sided paired Wilcoxon rank-sum test is used to test the difference of MCCs in 50 experiments between two deep learning methods among TLVar, Base-model and Self-model. The red dash line is the threshold of $-log_{10}(0.05)$.

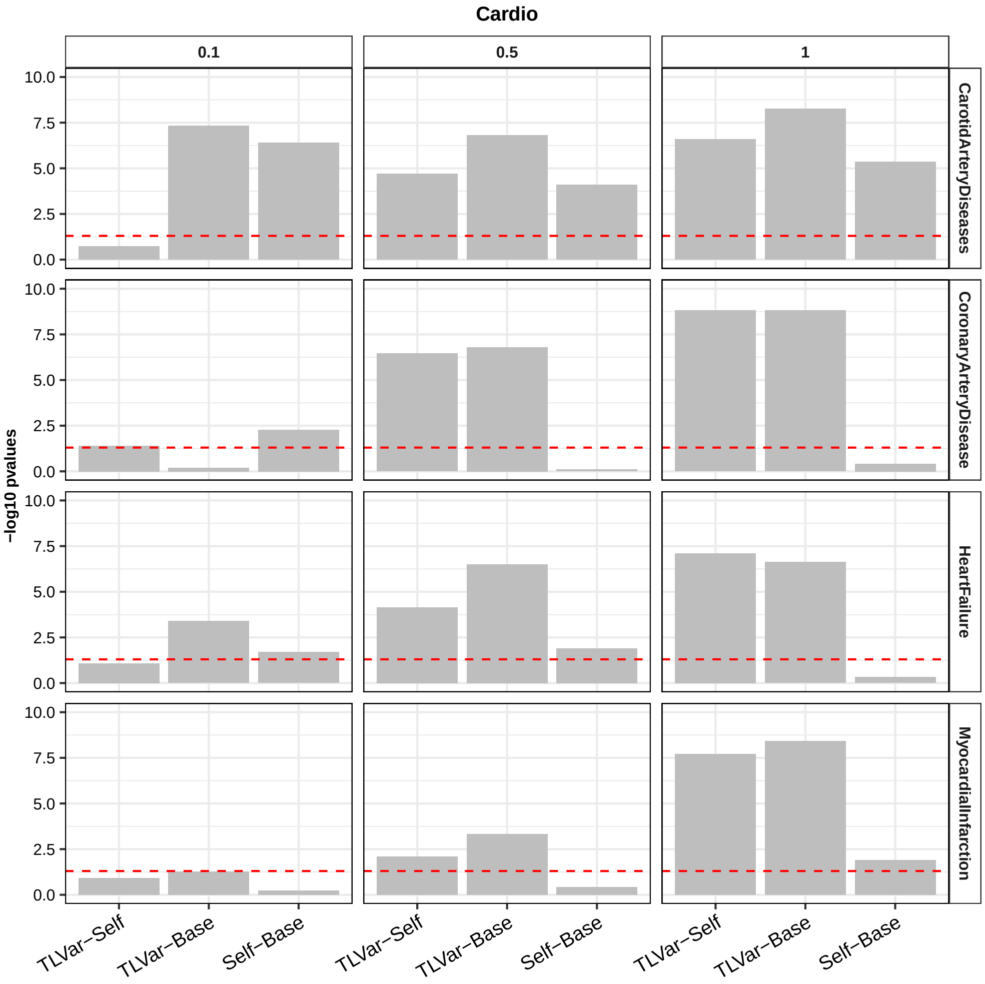


Figure S6: For 4 GWAS dataset in cardiovascular diseases, two-sided paired Wilcoxon rank-sum test is used to test the difference of AUCs in 50 experiments between two deep learning methods among TLVar, Base-model and Self-model. The red dash line is the threshold of $-log_{10}(0.05)$.

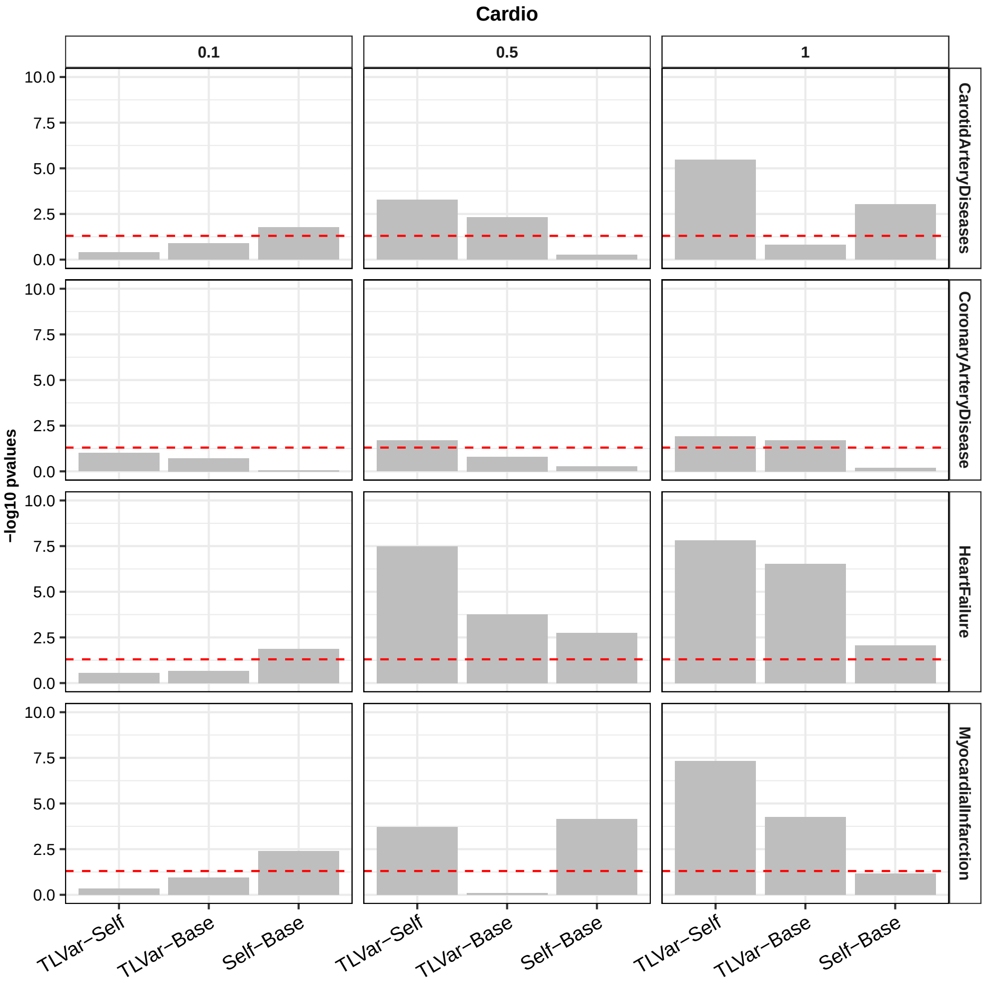


Figure S7: For 4 GWAS dataset in cardiovascular diseases, two-sided paired Wilcoxon rank-sum test is used to test the difference of MCCs in 50 experiments between two deep learning methods among TLVar, Base-model and Self-model. The red dash line is the threshold of $-log_{10}(0.05)$.

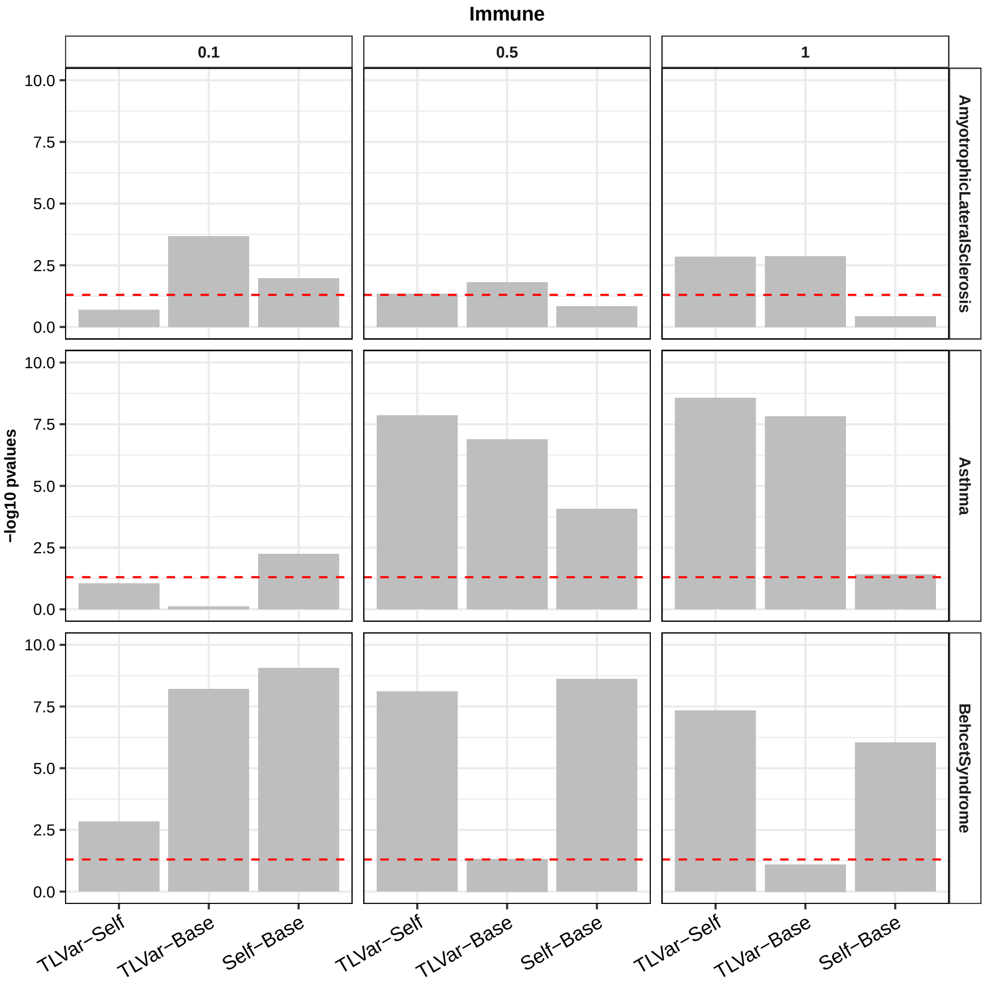


Figure S8: For 3 GWAS dataset in immune diseases, two-sided paired Wilcoxon rank-sum test is used to test the difference of AUCs in 50 experiments between two deep learning methods among TLVar, Base-model and Self-model. The red dash line is the threshold of $-log_{10}(0.05)$.

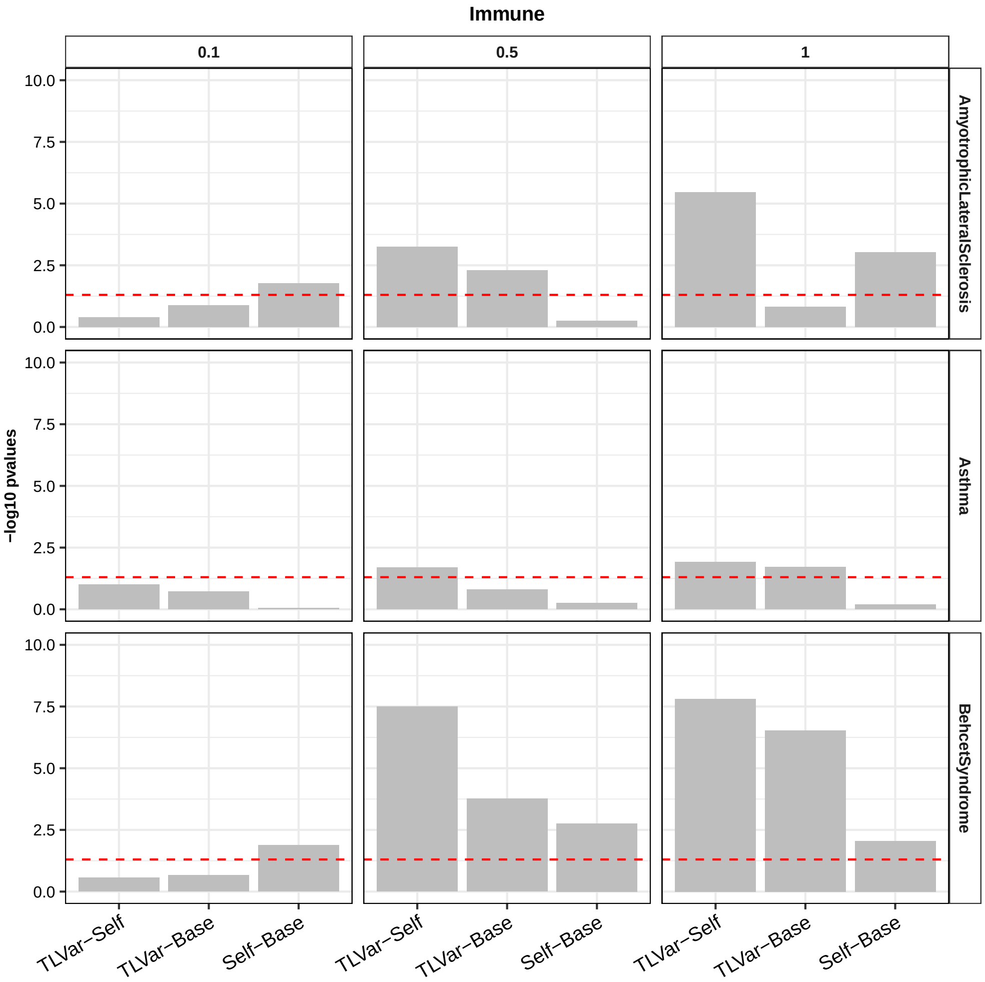


Figure S9: For 3 GWAS dataset in immune diseases, two-sided paired Wilcoxon rank-sum test is used to test the difference of MCCs in 50 experiments between two deep learning methods among TLVar, Base-model and Self-model. The red dash line is the threshold of $-log_{10}(0.05)$).

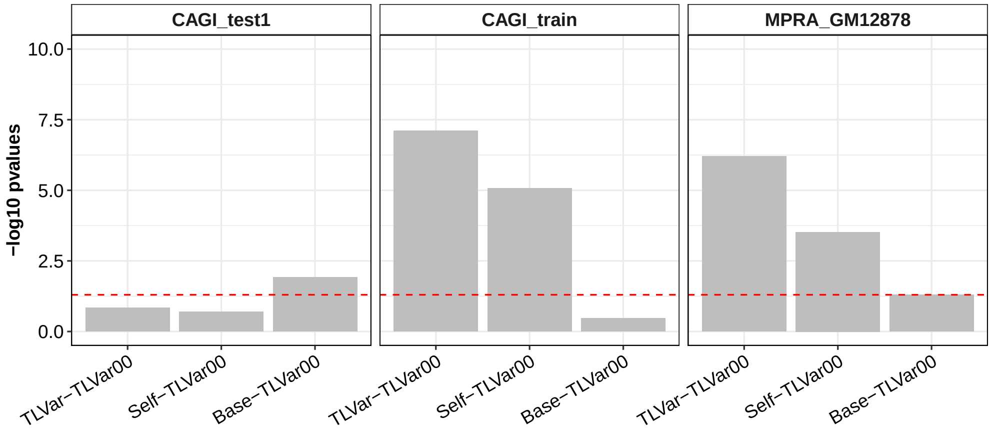


Figure S10: Deep learning method TLVar00, which is the same as TLVar except without convolutional layers not pretrained, is compared to abovementioned deep learning models in terms of AUC in three MPRA datasets. We randomly sample 20% MPRA regulatory variants as the independent testing set for all models and remaining 80% as the training set among which 20% is used as the validation set. To control the sampling bias, all experiments are repeated 50 times. two-sided paired Wilcoxon rank-sum test is used to test the difference of AUCs in 50 experiments between TLVar00 and each model among TLVar, Base-model and Self-model. The red dash line is the threshold of $-log_{10}(0.05)$.

### References


[1] Asa Ben-Hur and J Weston. Pyml-machine learning in python, 2010.
